# Disrupted neural adaptation in autism spectrum disorder

**DOI:** 10.1101/286880

**Authors:** Rachel Millin, Tamar Kolodny, Anastasia V. Flevaris, Alex M. Kale, Michael-Paul Schallmo, Jennifer Gerdts, Raphael A. Bernier, Scott O. Murray

**Affiliations:** Department of Psychology, University of Washington, Seattle WA 98195; Department of Psychiatry and Behavioral Sciences, University of Washington, Seattle WA 98195

## Abstract

There has been long-standing speculation that autism spectrum disorder (ASD) involves an increase in excitation relative to inhibition. However, there is little direct evidence of increased neural excitation in humans with ASD. Here we provide a potential explanation for this discrepancy: we show that increased neural excitation emerges only after repeated stimulation, manifesting as a deficit in neural adaptation. We measured fMRI responses induced by repeated audio-visual stimulation and button presses in early sensory-motor cortical areas. Across all cortical areas we show reduced adaptation in individuals with ASD compared to neurotypical individuals. The degree of adaptation is correlated between cortical areas and with button-press reaction times across subjects. These findings suggest that increased neural excitation in ASD, manifesting as dysregulated neural adaptation, is domain-general and behaviorally-relevant.

## Introduction

Autism spectrum disorder (ASD) is a behaviorally-defined, heterogeneous disorder with significant genotypic and phenotypic complexity. This complexity has made it challenging to identify the underlying neural mechanism(s) that are disrupted in the disorder. However, a unifying theme of numerous proposals is that the pathophysiology of autism is related to a pervasive increase in neural excitability^1–3^ – the likelihood of a neuron responding with an action potential to a given input signal. However, outside of cases in specific animal models of autism^4-6^ experimental evidence for increased neural excitation has been limited. Studies in humans across a variety of brain regions and measurement techniques have yielded equivocal results, with recent studies finding equivalent^7^, or even decreased^8^ neural response magnitudes in subjects with ASDs. We hypothesized that changes in neural excitation in ASD might manifest not in the initial transient neural response but, instead, in how neural responses adapt to repeated stimulation. This hypothesis was motivated by behavioral observations of reduced adaptation in ASD across diverse domains, including tactile stimulation^9–11^, face discrimination^12,13^, gaze direction^14^, numerosity^15^, audio-visual asynchrony^16^, and saccade amplitude^17^. To test this hypothesis, we used fMRI to characterize neural response amplitudes across time in individuals with ASD and neurotypical (NT) subjects.

## Results

We measured the fMRI response to repeated audio-visual stimulation and associated button press responses in early visual, auditory, and somato-motor cortical areas (Fig 1). We first compared the mean BOLD timecourses of subjects with ASD and NT subjects in a *regular* timing condition, in which stimuli were always presented 2 seconds apart within a 20 second block (Fig 1a). There was no significant difference in the initial fMRI response magnitude (4-8 seconds after the start of the block, after accounting for the hemodynamic delay) between groups in any cortical area examined (t(38)=-0.100, t(42)=0.199, and t(39)=-0.767 for auditory, visual, and motor, respectively; p=0.92, 0.91, and 0.45 as per two-tailed, two-sample t-tests). However, after this initial peak, the mean BOLD timecourse for the NT group was visibly reduced compared to that of the ASD group (Fig 1c). This difference between groups that develops after the initial response is consistent with increased neural excitability in ASD that manifests only after repeated stimulation, reflecting disrupted neural adaptation.

**Figure 1.**
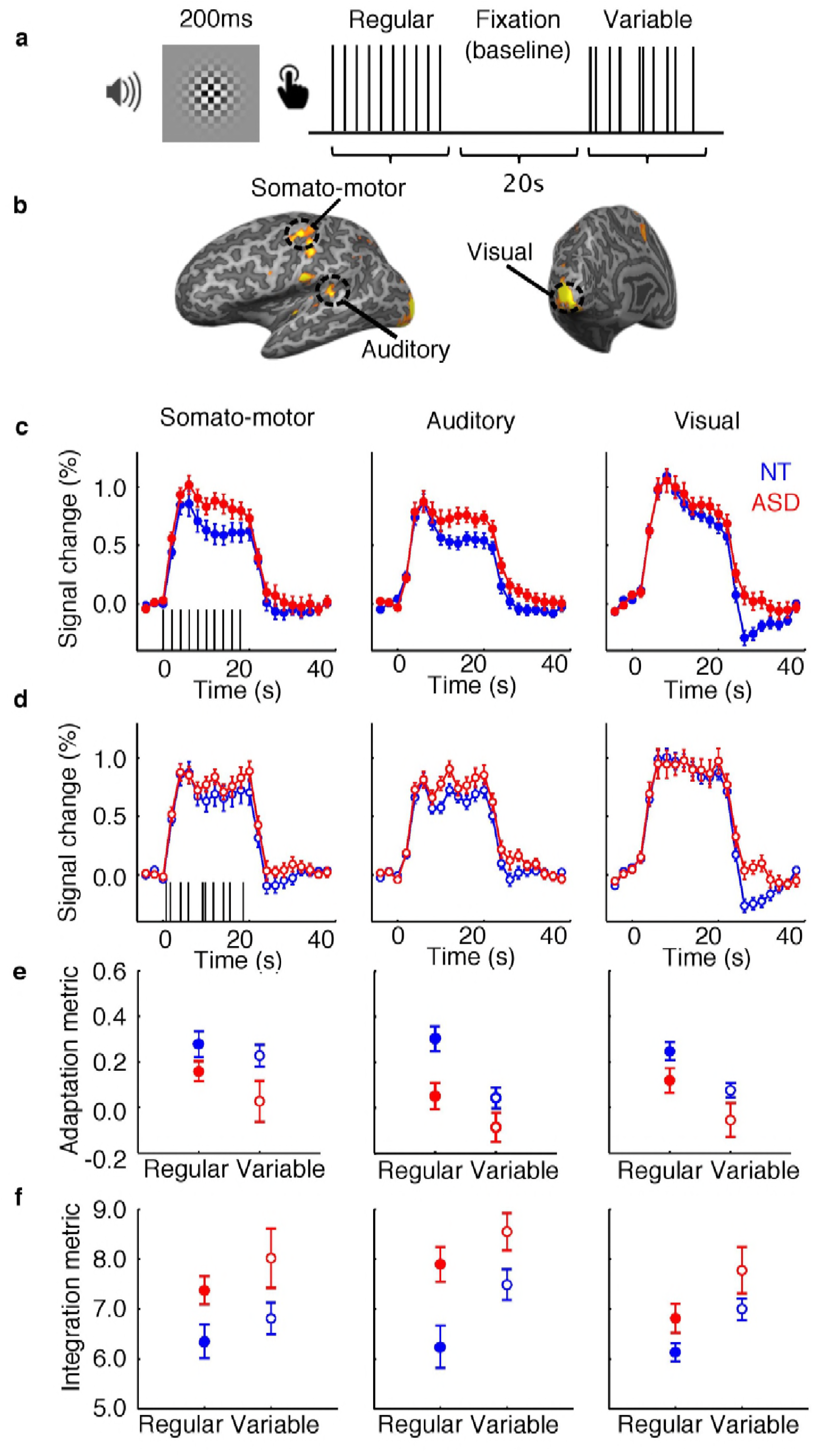
Task, ROI selection, and ROI timecourses. (a) The stimulus consisted of a checkerboard presented for 200ms accompanied for the duration by auditory white noise. Subjects were asked to respond with a button press as quickly as possible following the stimulus. Stimuli were presented with regular or variable inter-stimulus intervals in 20s blocks. The order of regular and variable blocks within the three runs was randomized, and this order was maintained across subjects. Stimulus blocks alternated with 20s of fixation. (b) ROIs in left and right visual, left and right auditory, and left or right (depending on handedness) somato-motor areas were selected based on activation to stimulus versus fixation blocks over the three experimental runs. ROI locations for an example subject are displayed. (c) Mean fMRI response timecourses in the regular timing blocks for NT (blue) and ASD (red) groups in somato-motor (left, N(NT)=24, N(ASD)=17), auditory (center, N(NT)=22, N(ASD)=18), and visual (right, N(NT)=24, N(ASD)=20) ROIs. (d) Same as (c), but for the variable condition. (e) Mean adaptation indices in somato-motor (left), auditory (center), and visual (right) ROIs for NT (blue) and ASD (red) groups in regular and variable conditions. (f) Same as (e), but for integration indices. Error bars indicate the standard error of the mean.

Next, we examined whether the differences between the NT and ASD groups depended on the statistical structure of the stimulus sequence. Specifically, by eliminating the regularity of the sequence we could assess whether the reduced adaptation depended on total stimulation history or was driven by temporal predictability. The stimuli in this *variable* condition occurred pseudorandomly with overall stimulation equal to that of the *regular* condition (Fig 1a). By visual inspection (Fig 1d), it is clear that the timecourses for both NT and ASD groups in the variable condition do not show the same reduction over time as in the regular condition: indeed, the timecourse of the fMRI signal is sensitive to the statistical structure of the stimulus timing. Interestingly, the separation between timecourses for NT and ASD groups was evident, though less prominent, when stimulus timing was unpredictable.

To quantify the degree of adaptation in the response timecourses, and the resulting overall total summed response between groups and across subjects, we calculated two metrics: an adaptation metric 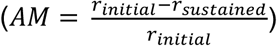 that reflects the signal decay relative to the initial response; and an integration metric 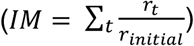 that reflects the total relative stimulus-evoked response for each subject in each condition (see Methods). Though the two metrics provide similar characterizations of the time courses they emphasize different components – the change in response over time vs. the cumulative response. A 3-way ANOVA on the AM (group x condition x ROI) showed main effects of group (F(1,31)= 9.82, p=0.0038), with lower AM for ASD subjects, and condition (F(1,31)=17.22, p=0.00024), with higher AM in the regular condition, but no interaction between the two (F(1,31)=0.20, p=0.66), or interaction between group and ROI (F(1.77,55)=0.14, p=0.85). The effect was mirrored by higher IM in the ASD group (group: F(1,31)=10.90, p=0.0024; condition: F(1,31)=10.17, p=0.0033; group x condition: F(1,31)=0.15, p=0.70; group x ROI: F(1.94,60.3)=0.85, p=0.43). In summary, we observed reduced adaptation and increased overall excitation in ASD compared to NT subjects, regardless of experimental condition.

The disrupted adaptation that we observed in ASD was present in both sensory and motor areas; further, the degree of adaptation was correlated between brain regions (Fig 2). Across all participants, we found a significant correlation between AM in auditory cortex and visual cortex (averaged across regular and variable conditions: Pearson’s r_36_=0.43, bootstrapped p=0.0082) auditory cortex and somato-motor cortex (r_33_=0.70, p<0.0001) and visual cortex and somato-motor cortex (r_36_=0.33, p=0.045). These correlations did not simply reflect reduced adaptation in the ASD compared to NT group in both areas; positive correlations were observed within both NT and ASD groups (Supplementary Material).

**Figure 2.**
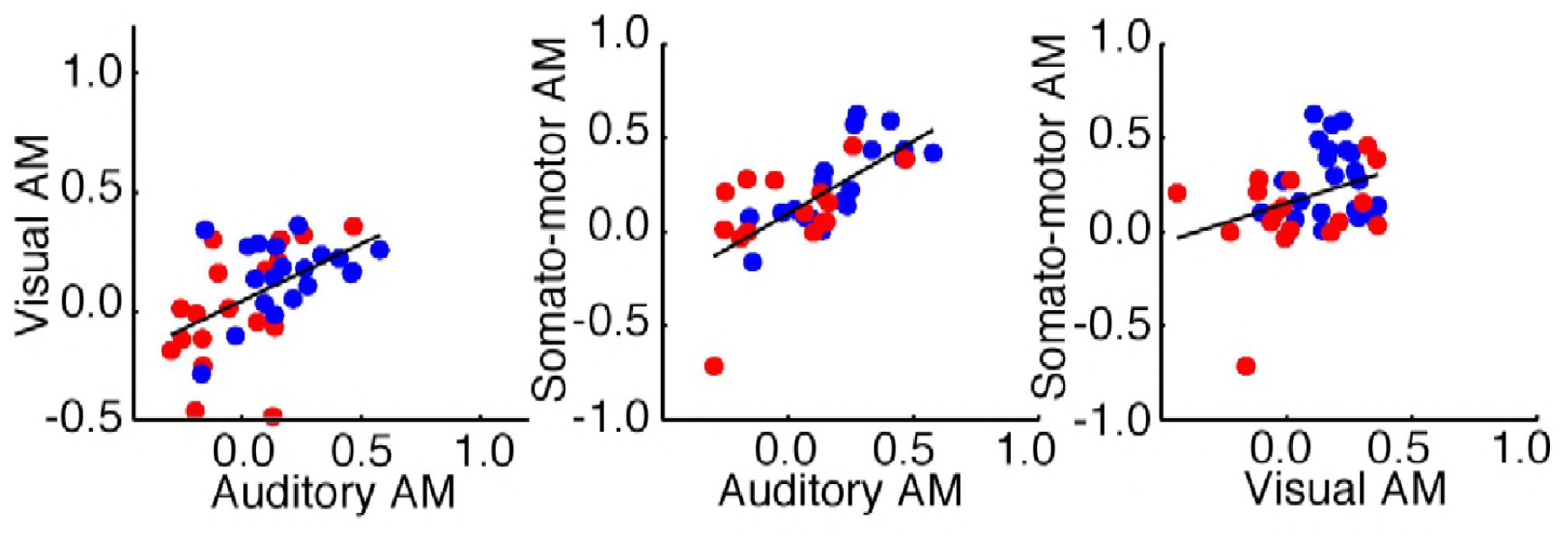
Relationship between AM across cortical areas. Correlation across subjects between mean AM across conditions in auditory and visual (left; N(NT)=20, N(ASD)=18), auditory and somato-motor (center; N(NT)=20, N(ASD)=15), and visual and somato-motor (right; N(NT)=21, N(ASD)=17) ROIs. Data for NT and ASD subjects are shown in blue and red, respectively. Black lines show linear fits to the data.

The divergence in BOLD response between groups reflects a difference in the neural response to repeated stimulation. However, it is possible that these differences could be compensated for by other processes in such a way that they do not influence behavior. To determine whether this effect does in fact have behavioral relevance, we examined the task performance of subjects while in the scanner. While ASD and NT subjects had similar reaction times to the first stimulus presentation in a block (2-way ANOVA, group x condition – group: F(1,46)=0.086, p=0.77; condition: F(1,46)=2.02, p=0.16; group x condition: F(1,46)=0.51, p=0.48), by the second presentation ASD subjects responded significantly more quickly to the stimulus; this effect persisted for the duration of the block (3-way ANOVA, group x condition x time – group: F(1,46)=4.30, p=0.044; time: F(3.42,157)=207, p<0.0001; group x time: F(3.42,157)=4.18, p=0.0049; group x condition x time: F(5.92,272)=1.12, p=0.35; Fig 3a). To test whether this performance difference was associated with the adaptation difference, we examined the correlation between reaction time (RT) and AM. We found that, across all subjects, faster RT was associated with lower AM (Supplementary Material). This correlation was strongest (r_39_=0.48, p=0.0014), and present within both NT (r_22_=0.40, p=0.05) and ASD groups (r_15_=0.51, p=0.036), in the regular condition in the somato-motor ROI (Fig 3b).

**Figure 3.**
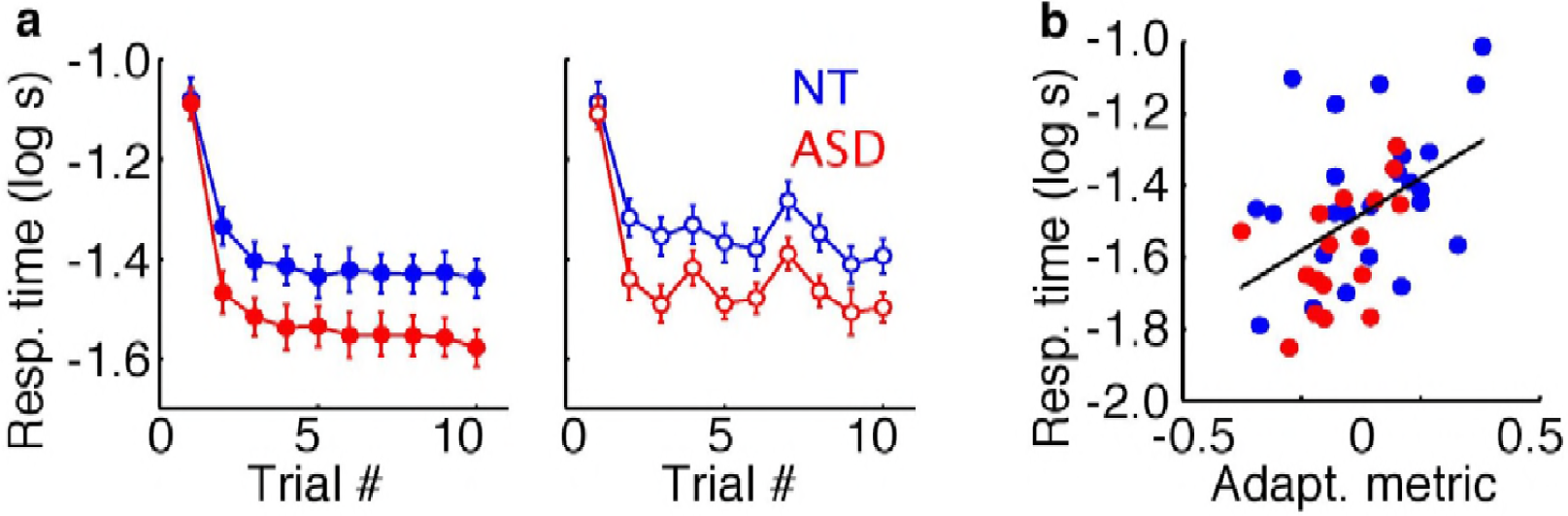
Reaction time data (log-transformed) for NT (blue) and ASD (red) subjects. (a) Mean time between stimulus onset and button press (RT) as a function of trial within a block, averaged across blocks, in regular (left) and variable (right) timing conditions for 27 NT and 20 ASD subjects. Error bars indicate the standard error of the mean. (b) Correlation in the regular condition between AM in the somato-motor ROI and mean RT across all but the first trial for the 24 NT and 17 ASD subjects with usable data for this ROI. Black line shows a linear fit to the data.

To rule out potential alternative explanations for the reduced adaptation observed in our ASD group, we examined NT and ASD subgroups matched for gender, and found the same pattern of results (Supplementary Fig 1). We also analyzed head motion, eye movement behavior, and task engagement as indexed by behavioral performance. For all three measures we found that there was either no difference between subject groups (behavioral performance: first RT (Fig 3) and false alarms; eye movement behavior: time spent fixating, distance of center of gaze from fixation mark) or that there was no statistical contribution to the effect (head motion: change in head position between successive TRs; Supplementary Fig 2-5).

## Discussion

Overall, our results support the hypothesis that deficits in a fundamental neural regulatory process, adaptation, result in overall increased neural excitation in ASD during sustained task performance. We observed reduced adaptation in varying degrees across auditory, visual, and somato-motor areas. The degree of adaptation was correlated between cortical areas across subjects, consistent with this difference being a domain-general individual trait. While previous research has shown decreased repetition suppression in ASD to socially-relevant stimuli such as faces^18^ the degree to which repetition suppression is an index of neural adaptation or priming is an open question^19,20^. In addition, subjects with ASD show less reduction in fMRI response than NT subjects to aversive stimulation, measured in successive blocks of presentation^21^; without a manipulation of stimulus properties it is unclear whether this effect reflects neural or hemodynamic properties or whether it relies on the same processes as the rapidly-appearing differences observed here^22,23^.

While we observed differences in adaptation in primary sensory cortical areas that are involved in the earliest stages of processing of sensory stimuli, it is possible that these differences emerge due to modulation via top-down or feedback signals. For example, accuracy of stimulus expectation, which has been proposed to contribute to repetition suppression in visual areas^19^, could differ between NT and ASD groups. In fact, theoretical work has suggested that many of the core symptoms of ASD, including the failure to adapt or habituate, can be framed as an inability to form predictions^24^ and incorporate prior experiences^25^. Here, differences in adaption did not appear sensitive to the statistical regularity of the stimulation sequence. A feedback-based explanation also seems to be at least partially at odds with our RT data: subjects with ASD perform better than NT subjects with repetition in both regular and variable conditions, and their advantage is similar regardless of stimulus predictability. Considering the correlation between AM and RT, a potentially more parsimonious explanation of our results is that differences in local processing lead to reduced adaptation in ASD, which results in an advantage in the button press task. While the present study provides evidence for increased excitation in individuals with ASD that emerges over time due to reduced adaptation, future work is necessary to delineate the relative contributions of locally-mediated versus feedback contributions to disrupted adaptation.

## Materials and Methods

### Subjects

24 subjects with autism spectrum disorder (ASD; 16 males) and 29 neurotypical (NT) subjects (14 males) participated in the experiment. 4 of these subjects (2 males with ASD, 1 NT male, and 1 NT female) were excluded prior to analysis due to excessive head motion (consistent head movements of >1 mm). 2 additional subjects (1 male and 1 female with ASD) had no usable data as per the data cleaning procedure described below, resulting in 27 NT subjects (13 male) and 20 subjects with ASD (13 male) included in subsequent analysis. This sample size is similar to those reported in relevant, methodologically-similar recent studies^7,26^. All subjects had normal IQ (WASI-II Full Scale IQ of at least 80), and normal or corrected-to-normal vision. Groups were age- and IQ-matched (mean IQ of subjects with autism: 114; NT subjects: 112; t(45)=0.39, p=0.70; mean age of subjects with autism: 23 years; NT subjects: 24 years; t(45)=-0.96, p=0.34). All subjects provided written informed consent to participate. The Institutional Review Board of the University of Washington (UW) approved the research protocol. Subjects with ASD met diagnostic criteria for ASD on the Autism Diagnostic Interview-Revised (ADI-R^27^), the Autism Diagnostic Observation Schedule—2^nd^ Edition (ADOS-2^28^) and according to expert clinical judgment using DSM-5^29^ criteria. All group analyses were repeated with a subset of subjects (13 ASD, 8 males, and 13 NT, 4 males), matched for head motion in the scanner to control for its potential impact on fMRI signal, and a subset matched for gender (12 ASD, 6 males, and 12 NT, 6 males, see Supplementary Material).

### MRI Acquisition

Scans were acquired with a Philips Achieva 3T MRI system. A T1-weighted structural scan was acquired at the beginning of the scan session, followed by three functional gradient-echo EPI scans with axial orientation (30 slices with 3 mm inplane resolution and 0.5 mm gap, 2 s TR, 25 ms TE, 79° flip angle, A-P phase-encode direction). A single TR EPI scan with opposite phase-encoding direction (P-A), but otherwise identical to those above, was acquired for use in correcting geometric distortions. Each subject underwent a single scanning session, lasting approximately one hour (the scan session also included acquisition of spectroscopy data for a separate experiment). When possible, subjects’ eyes were tracked during scanning using an Eyelink 1000 Plus eyetracker, sampling at 1000 Hz. Due to the challenges of eyetracking in the scanner, we were able to collect data for 55 % of subjects (N(NT) = 17, N(ASD) = 9).

### Stimuli

Stimuli were presented using Presentation software running on a Windows XP computer. Images were projected onto a screen behind the subject’s head via either an Epson Powerlite 7250 or an Eiki LCXL100A projector (following a hardware failure), both operating at 60 Hz, and with linearized luminance profiles. Subjects viewed the projected images using a mirror positioned above their eyes, for an effective viewing distance of 66 cm. Sound was delivered at 44.1 kHz using MRI compatible earbuds (S14, Sensimetrics). Subjects wore protective ear muffs over the earbuds to attenuate acoustic noise from the scanner. Before scanning, subjects verified that the auditory stimulus was presented at an audible and comfortable volume.

The stimulus consisted of a Gaussian-windowed (with FWHM of 2 deg. visual angle), full-contrast checkerboard (check size of 0.4 deg) image presented on a uniform background accompanied by audio white noise. Visual and auditory stimuli were presented simultaneously for 200 ms in a blocked design, with the stimulus presented 10 times within each stimulus block. Blocks were of two types: regular and variable. In regular blocks, the stimuli were separated by 1800 ms of rest, resulting in a stimulus presentation every 2 seconds. In variable blocks, the inter-stimulus interval was a random value drawn from a uniform distribution bounded by 800 ms and 2800 ms. Stimulus blocks alternated with rest blocks. A fixation cross appeared at the center of the display whenever the stimulus was off. Each subject completed three runs of 8 stimulus blocks and 9 rest blocks each. The protocols (stimulus timing) for these runs were identical across all subjects, but the variable blocks differed across the 3 fMRI scans. Subjects were instructed to use the index finger of their dominant hand to press a button as quickly as possible following the appearance of each stimulus. Subjects completed one practice run during a mock scan session prior to scanning.

### MRI data preprocessing

Data was preprocessed using BrainVoyager (Brain Innovation, Maastricht, The Netherlands) software. EPI data was motion-corrected, corrected for distortion due to magnetic field inhomogeneities, high-pass filtered (cutoff = 2 cycles/scan), and coregistered to the AC-PC-aligned T1 structural scan.

To identify regions of interest corresponding to early visual, auditory, and somato-motor cortical areas, statistical activation maps were determined from z-transformed data from all three functional runs. Variable and standard conditions were treated as a single condition (versus rest) to generate a boxcar predictor, which was then convolved with a double gamma HRF. This predictor plus the 3 translation and 3 rotation parameters obtained during motion correction were used as the design matrix in a general linear model that was fit to the timecourse of each voxel. The resulting activation maps from the t-statistic for the model fit were initially thresholded at p < 0.05 (Bonferroni corrected). ROIs were selected manually from the most significant areas of activation near visual (left and right), auditory (left and right), and somato-motor (contralateral to the dominant hand) cortices, yielding 5 ROIs for each subject. If multiple activation clusters were present, ROIs were selected according to the following criteria. For the early visual ROI, the cluster nearest the occipital pole, in line with the calcarine sulcus, was selected. For the somato-motor ROI, the cluster closest to the hand knob, the omega-like structure in the pre-central gyrus^30^, was chosen. The anatomical location of auditory activation was variable across subjects; the most significantly activated cluster on the superior temporal lobe was selected. If no obvious cluster of voxels was present even after the threshold was lowered, the ROI was excluded. The 20 most significantly activated voxels in the cluster defined the final ROI in each region. For some ROIs in some subjects, fewer than 20 voxels met the threshold criteria; in these cases, the threshold was relaxed until 20 voxels could be selected. If no obvious cluster of voxels was present after the threshold was lowered, the ROI was excluded. This resulted in 46 subjects with data for the auditory ROI, 48 for the visual ROI, and 46 for the motor ROI.

### fMRI data analysis

Average timecourses across the 20 voxels in each ROI were determined for each run. Percent-transformed timecourses were then calculated for each block type by first extracting the data in an epoch spanning from 2 TRs (4 seconds) prior to a stimulus block to 10 TRs (20 seconds) after its end. Each such timecourse was then normalized by subtracting and dividing by the first 3 TRs (these TRs were treated as a baseline), and multiplying by 100. The resulting block timecourses were then averaged over all blocks of the same condition, yielding one such timecourse for variable timing blocks and another for regular blocks. Blocks that did not meet criteria for head motion and task performance were excluded prior to averaging. Data for a given block was excluded due to head motion if the subject’s head moved more than 0.9 mm between two successive TRs (frame-wise displacement > 0.9 mm^31^) up to and including 8 TRs before or 1 TR after the stimulus block. A block was excluded on the basis of task performance if it contained any misses (failure to press the button after a stimulus appearance) or more than one false alarm (more than one button press after appearance of a stimulus). If more than half the blocks of either condition were excluded, or a subject had more than 7 behavioral errors in a run, the entire run was excluded. If 2 out of 3 runs were excluded, all data for the subject were excluded. 7 out of 147 total runs (1 NT and 6 ASD) and 1 complete subject (ASD) were excluded using this procedure. Noisy data were further removed by excluding the resulting timecourses for which less than 50% of the power (as per discrete Fourier analysis of the ROI timecourse) was at 1 cycle per duration of the extracted event-related timecourse (44s). This resulted in the exclusion of 1 additional ASD subject, and 14 of the remaining subjects retaining data for only one or two of the three ROIs. With cleaning, data was retained for 22 NT and 18 ASD subjects in the auditory ROI, 24 and 20 in the visual ROI, and 24 and 17 in the somato-motor ROI. An adaptation metric (AM) and integration metric (IM) were calculated for each subject in each ROI from the resulting set of timecourses. AM was calculated as the initial response to the stimulus blocks (defined as 4 to 6 seconds after block onset for auditory and somato-motor cortex; 6 to 8 seconds after block onset for visual cortex, where the hemodynamic response had a longer delay) minus the response 12 to 20 seconds after the start of the block, normalized by the initial response 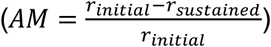. IM was calculated as the sum across timepoints in this window normalized by the initial response 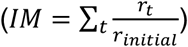.

### Statistics

All statistical analyses were performed in Matlab 2013b or SPSS 19. We assume that our data are normally distributed and use standard parametric tests to determine statistical significance. 2-tailed t-tests were used to test for differences between ASD and NT groups, and correlation was quantified with Pearson’s r. Significance of correlations was determined using a bootstrap procedure: correlation coefficient was recalculated 10000 times on data resampled with replacement independently for each variable to identify a percentile for the correlation coefficient. For ANOVAs, deviations from sphericity were corrected for using the Greenhouse-Geisser procedure. In the case of RTs, which typically have an ex-Gaussian distribution, we perform averaging and statistics on the log-transformed values.

## Supplementary Data

### AM is correlated between cortical areas within NT and ASD groups

To determine whether the correlation of AM between regions was driven only by group differences in AM across ROIs, we examined each group separately. In both groups, AM was positively correlated between most ROI pairs (auditory and visual – NT: r=0.34, p=0.14, ASD: r=0.52, p=0.029; auditory and somato-motor – NT: r=0.77, p=0.0002, ASD: r=0.55, p=0.029; visual and somato-motor – NT: r=0.037, p=0.87, ASD: r=0.29, p=0.27).

### AM is positively correlated with repeated button press RT

In all of the ROIs examined, AM was positively correlated with the mean RT over the stimulus block (excluding the first response in a block) in regular (auditory: r=0.43, p=0.0069; visual: r=0.23, p=0.13; motor: r=0.48, p=0.0014) and variable (auditory: r=0.32, p=0.045; visual: r=0.15, p=0.32; motor: r=0.33, p=0.035) conditions. In most cases, correlations were also positive within NT (regular - auditory: r=0.38, p=0.08; visual: r=0.11, p=0.62; motor: r=0.40, p=0.052; variable - auditory: r=0.11, p=0.61; visual: r=-0.17, p=0.43; motor: r=0.24, p=0.26) and ASD (regular – auditory: r=0.16, p=0.52; visual: r=0.10, p=0.67; motor: r=0.51, p=0.036; variable – auditory: r=0.43, p=0.071; visual: r=0.18, p=0.45; motor: r=0.30, p=0.24) groups.

### Differences in gender balance, head motion, eye movements, or task engagement between NT and ASD groups do not explain reduced adaptation in ASD

In order to rule out the contribution of nuisance factors to the results, we performed several supplementary analyses. First, we determined that the results were not driven by the different numbers of males and females in the NT and ASD groups (the ASD group had a higher proportion of male participants). We re-examined the AM and IM in a smaller group of subjects, with equal numbers of male and female subjects (6 each) in NT and ASD groups. All subjects included in this analysis had usable data for all ROIs. The pattern of results in this group was the same as that of the full group, with lower average AM and higher average IM in the ASD compared to NT group (Supplementary Fig 1). 3-way ANOVA (group x condition x ROI) on AM showed main effects of group (F(1,22)=4.85, p=0.039) and condition (F(1,22)=13.03, p=0.0016) but no interaction between the two (F(1,22)=0.68, p=0.42), or interaction between group and ROI (F(1.73,38.09)=0.37, p=0.66). Similarly, 3-way ANOVA (group x condition x ROI) on IM showed main effects of group (F(1,22)=5.95, p=0.023) and condition (F(1,22)=9.35, p=0.0058) but no interaction between the two (F(1,22)=0.16, p=0.69), or interaction between group and ROI (F(1.79,39.5)=1.27, p=0.29).

**Supplementary Figure 1.**
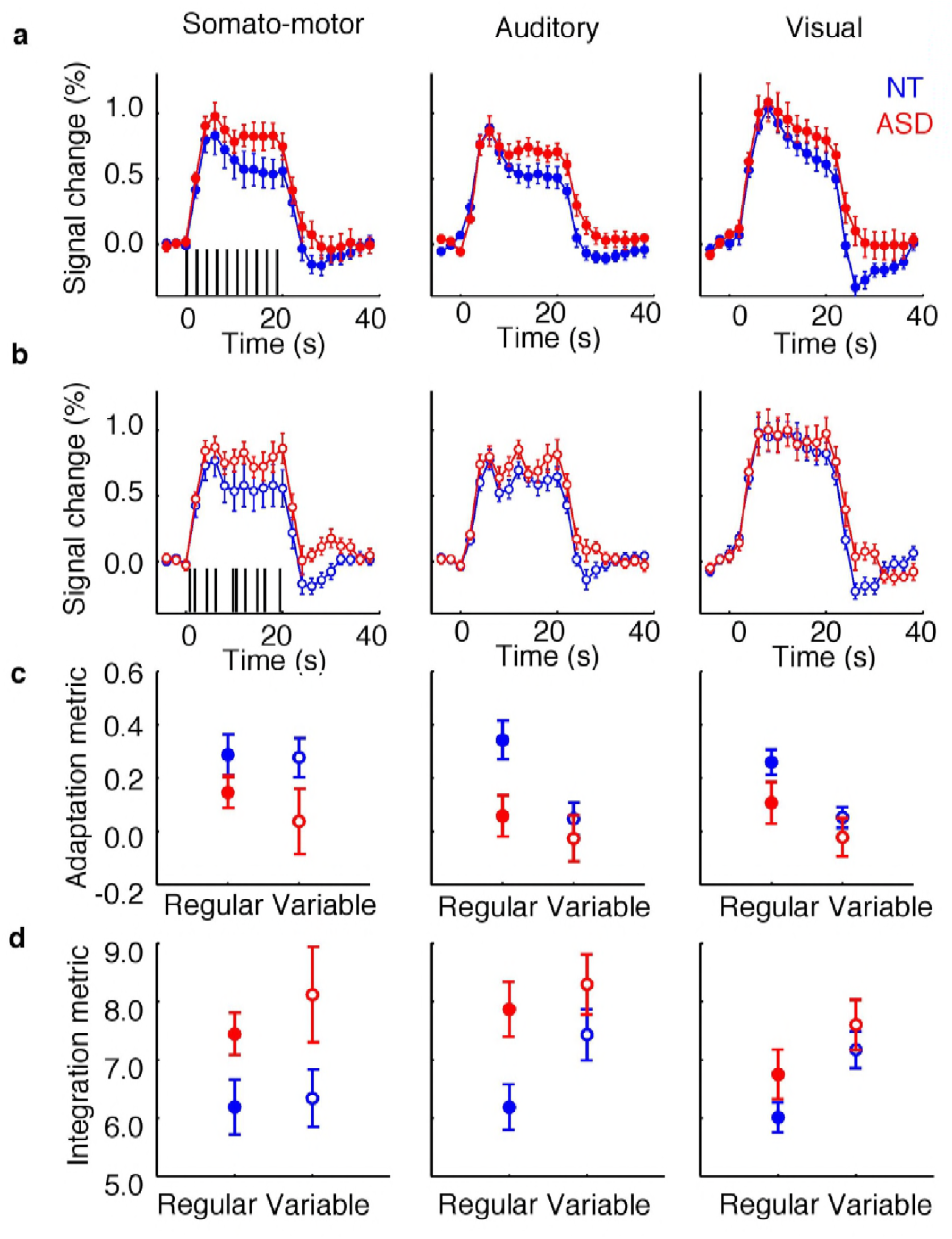
fMRI results for the 12 NT (blue) and 12 ASD (red) subjects matched by gender. (a) Mean fMRI response timecourses in the regular timing in somato-motor (left) auditory (center), and visual (right), ROIs. (c) Mean adaptation indices in somato-motor (left), auditory (center), and visual (right) ROIs for NT and ASD groups in regular and variable conditions. (d) Same as (c), but for integration indices. Error bars indicate the standard error of the mean.

Results were also not due to differences in head motion between NT and ASD groups, though differences were present. To quantify subjects’ head motion, we calculated frame-wise displacement (FWD, the difference in head position between adjacent TRs^1^) for all TRs used in our analyses. As a group, subjects with ASD moved their heads significantly more than NT subjects (mean FWD: t(46)=-2.72, p<0.01; FWD standard deviation: t(46)=-2.10, p<0.05; Supplementary Fig 2a). However, the differences between NT and ASD groups in AM (F(1,30)=5.60, p=0.010) and IM (F(1,30)=8.44, p=0.007) remained after adding mean FWD as a covariate in the 3-way ANOVA (group x condition x ROI). To further rule out head motion as an explanation for our results, we analyzed AM and IM in a subgroup of 13 NT and 13 ASD subjects matched for mean FWD (Supplementary Fig 2b), and found a similar pattern of results as in the larger group (Supplementary Fig 3). In this smaller group, AM differed significantly between ASD and NT groups (3-way ANOVA, group x condition x ROI, F(1,24)=6.93, p=0.015) and between conditions (F(1,24)=24.9, p=0.000043). This result was mirrored in the IM (group: F(1,24)=8.99, p=0.0062; condition: F(1,24)=12.33, p=0.0018).

**Supplementary Figure 2.**
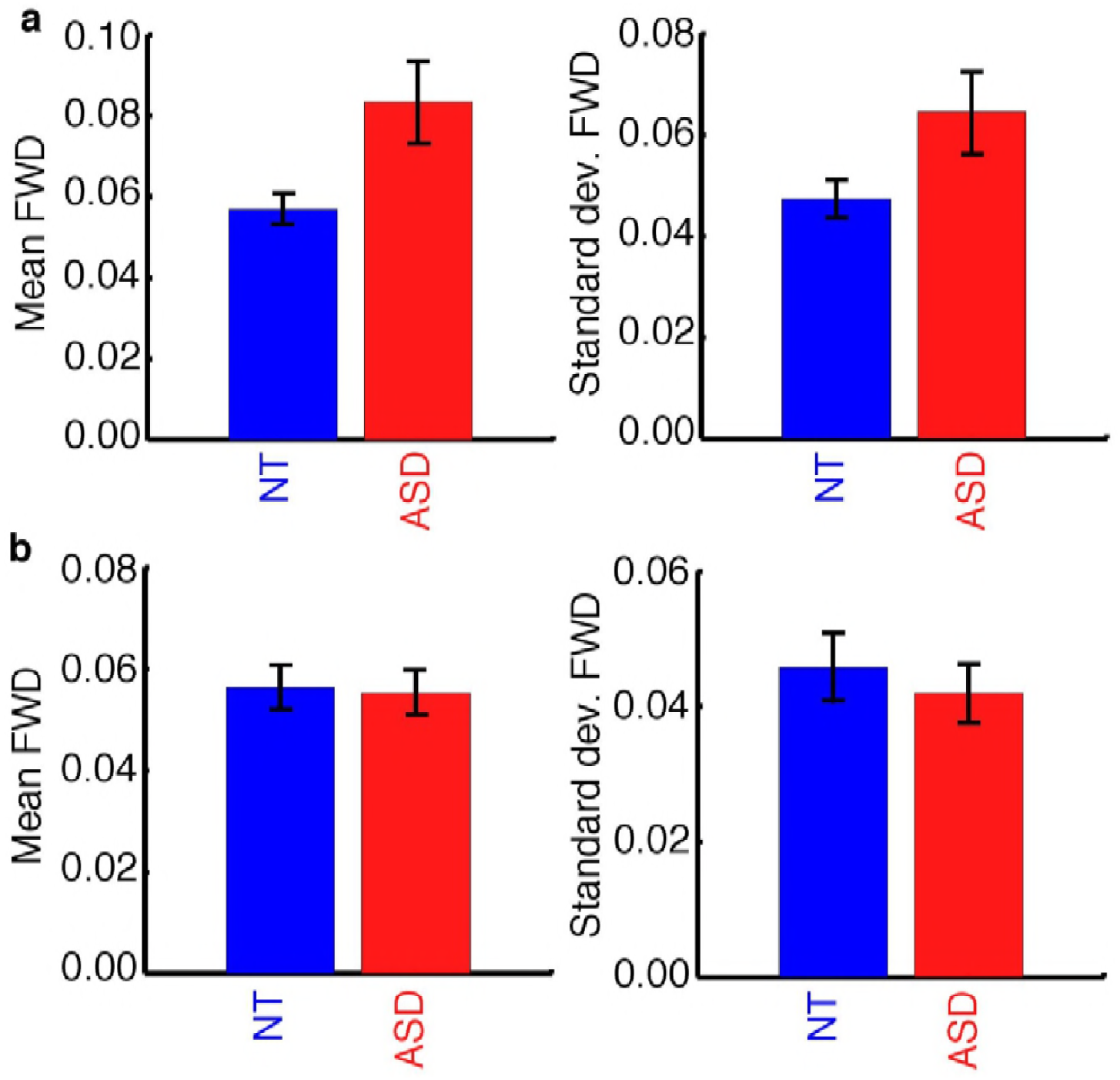
Head motion data for NT (blue) and ASD (red) subjects. (a) Mean (left) and standard deviation (right) of FWD displacement in the full group of 27 NT and 20 ASD subjects. (b) Same as (a) for the 13 NT and 13 ASD subjects matched for mean FWD.

**Supplementary Figure 3.**
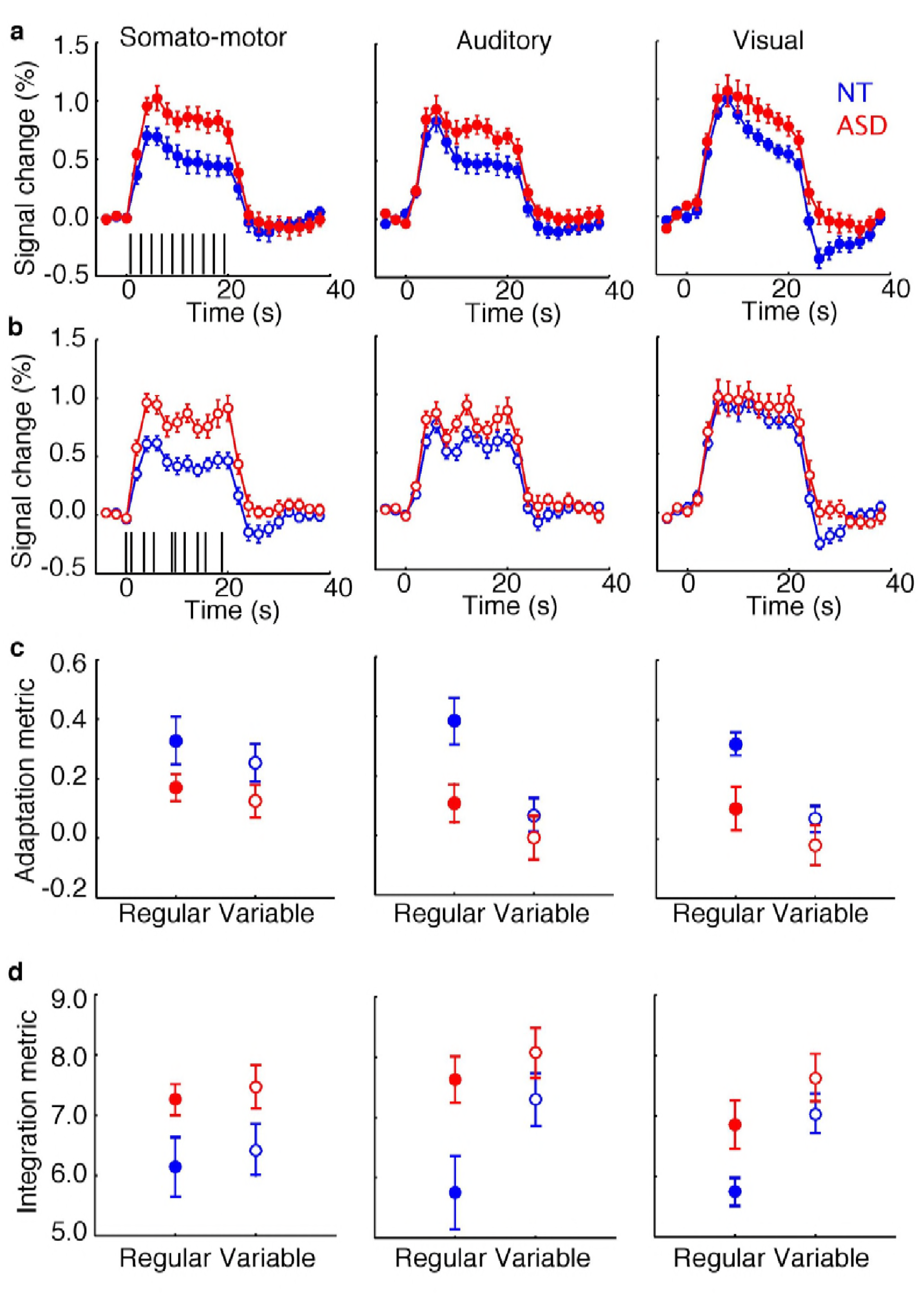
fMRI results for the 13 NT (blue) and 13 ASD (red) subjects matched by mean FWD. (a) Mean fMRI response timecourses in the regular timing in somato-motor (left), auditory (center), and visual (right) ROIs. (c) Mean adaptation indices in somato-motor (left), auditory (center), and visual (right) ROIs for NT and ASD groups in regular and variable conditions. (d) Same as (c), but for integration indices. Error bars indicate the standard error of the mean.

Group differences also cannot be explained by differences in eye movement behavior in the subjects whose eyes were successfully tracked (17 NT and 9 ASD in the full group, 7 NT and 9 ASD in the group matched by head movement). We found no difference in the proportion of time spent fixating (t(24)=-0.62, p=0.54; t(14)=-0.93, p=0.37), or the mean (t(24)=-1.18, p=0.25; t(14)=-0.90, p=0.38) or standard deviation (t(24)=-1.20, p=0.24; t(14)=-0.70, p=0.49) of the distance of eye position from the fixation mark in the full group or group matched by mean FWD, respectively (Supplementary Fig 4).

**Supplementary Figure 4.**
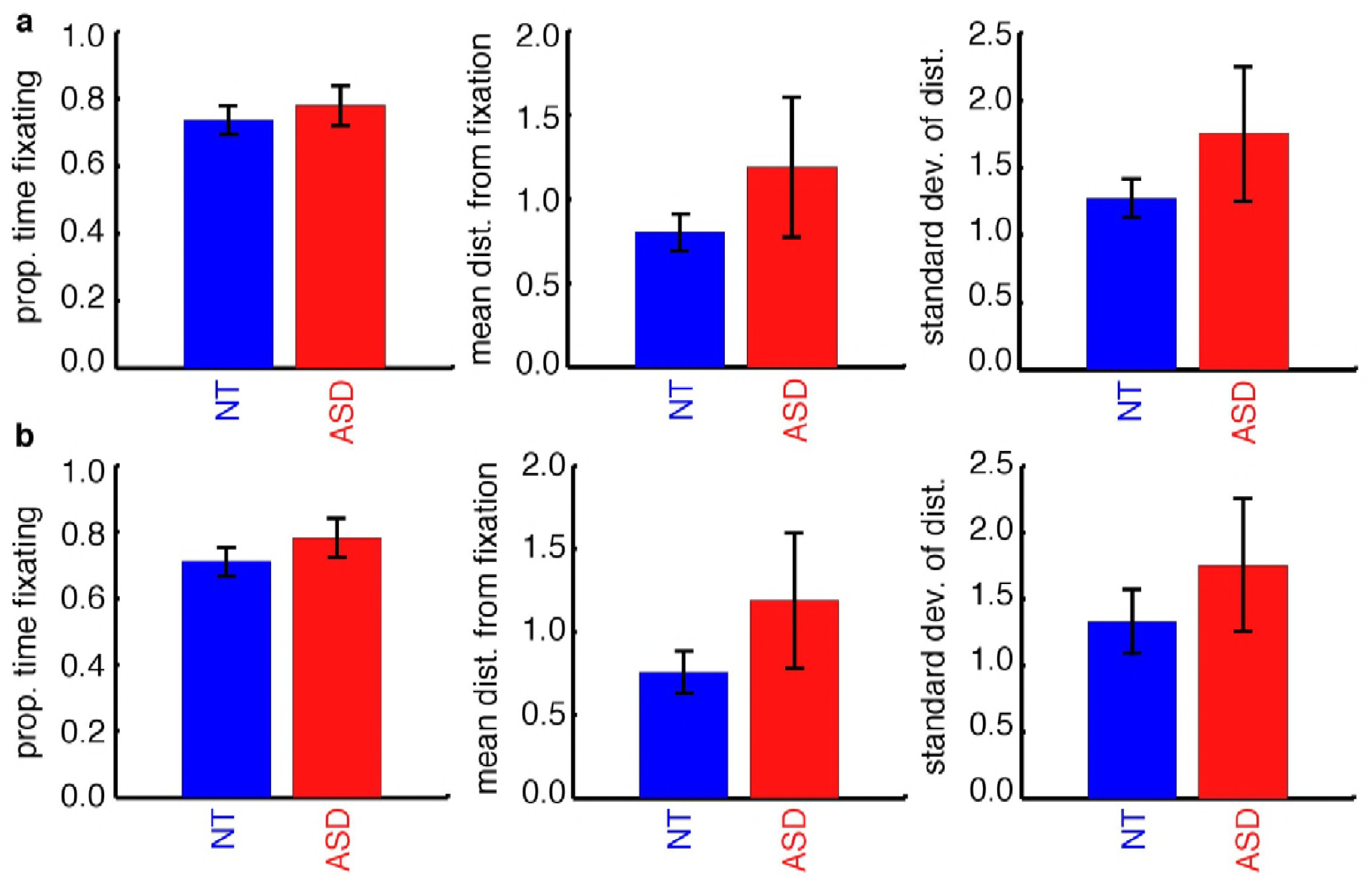
Eye movement results for NT (blue) and ASD (red) subjects. (a) Proportion of time spent fixating (left), mean distance of center of gaze from fixation (center), and standard deviation of the distance from fixation (left) in the full group of subjects with eye-tracking data (17 NT and 9 ASD). (b) Same as A but for subjects matched by mean FWD. (a) and (b) appear very similar because ASD subjects are the same in both analyses, and 7 NT subjects are included in both analyses. Error bars indicate the standard error of the mean across subjects.

Lastly, aside from the shorter RTs observed in the ASD group with repeated stimulus presentation, we did not find evidence for differences between the groups’ engagement in the task during blocks used in our analyses. After discarding fMRI and behavioral data based on head motion and performance (see Methods), there is no difference between NT and ASD groups in the number of false alarms (2-way ANOVA, group x condition, F(1,46)=0.71, p=0.13; F(1,24)=0.26, p=0.62), or the RT for the first stimulus in a block (F(1,46)=0.26, p=0.61; F(1,24)=2.05, p=0.17) for either the full group of subjects or the matched group, respectively (Supplementary Fig 5 and Fig 3).

**Supplementary Figure 5.**
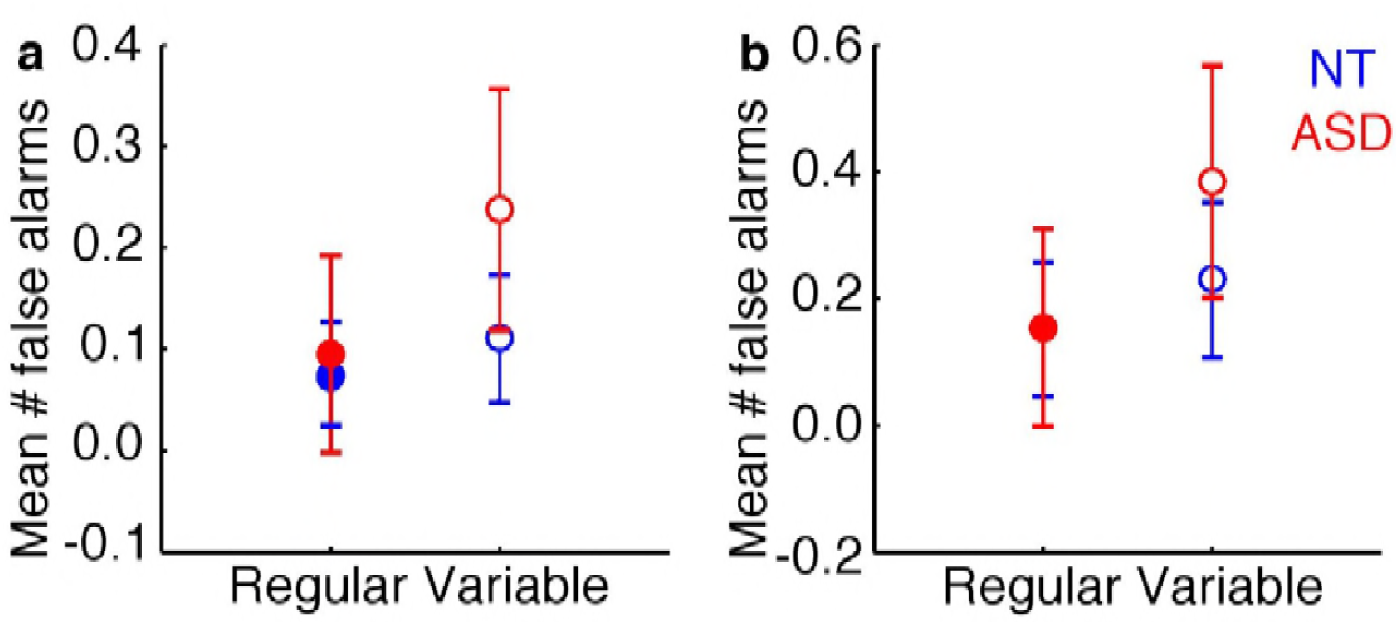
Button press results for NT (blue) and ASD (red) subjects. (a) Mean number of false alarms per stimulus block in the full group of subjects. (b) Same as (a) but for subjects matched by mean FWD. Error bars indicate the standard error of the mean across subjects.

